# Manual validation finds only ultra-long long-read sequencing enables faithful, population-level structural variant calling in *Drosophila melanogaster* euchromatin

**DOI:** 10.1101/2025.04.21.649852

**Authors:** James A. Hemker, Hannah R. Gellert, Jessica A. Smiley-Rhodes, Bernard Y. Kim, Dmitri A. Petrov

## Abstract

The increasing accessibility of long-read sequencing and the rapid development of automated variant callers are promoting the generation of population-level structural variation data. However, the effect of the length of long-reads on automated variant callers is not well understood, especially for non-human species. Here we show that only ultra-long long-reads, with read N50s greater than 50kb, are capable of accurately calling structural variants of any size in *Drosophila melanogaster* euchromatin. We used Oxford Nanopore Technologies to long-read sequence eight, inbred *D. melanogaster* strains to extremely high coverage (mean 238***×***), and we then downsampled the reads to create read pools of different length distributions. We assembled genomes from these different read-length pools and used both read-based and assembly-based structural variant callers to call variants in each strain before merging the calls into population-level datasets. We manually validated over 2,300 putative structural variants to assess the accuracy of the variant calls across the different read-length distributions and to determine the cause and rates of false positive errors. We found that more than half of all structural-variant-calling errors stem from misaligned reads that contain mobile elements or are located in repetitive and complex regions. Overall, our results show that long reads need to be at least three times longer than the repetitive and mobile elements found in the genome in order to accurately call structural variants at the population level.

## Introduction

Structural variants are genomic rearrangements such as insertions, deletions, duplications, and inversions that are often defined as greater than 50bp (Mahmoud et al. 2019; Ahsan et al. 2023). Historically, single-nucleotide polymorphisms (SNPs) have been the focus of genomic diversity metrics and studies, but further research has shown that structural variants affect a larger proportion of the genome than SNPs (Pang et al. 2010; Durbin et al. 2010; Catanach et al. 2019). Recent population genetics studies suggest that structural variants face stronger negative selection than SNPs (Chakraborty et al. 2019; Hämälä et al. 2021), and structural variation underpins many severe human diseases. Repeat expansions are implicated in both Parkinson’s disease (Alvarez Jerez et al. 2024) and Huntington’s disease (Walker 2007), and structural variants are the leading class of driver mutations in cancer, as they can fuse or disrupt oncogenes and repurpose regulatory elements for dysregulated gene expression (Cosenza et al. 2022). At the same time, structural variants have played critical roles as large-effect adaptive mutations across the tree of life (Wellenreuther et al. 2019; Kapun and Flatt 2019; Gorkovskiy and Verstrepen 2021; Hämälä et al. 2021; Shi et al. 2023; Ferguson et al. 2024; Dodge et al. 2024). A tissue-specific-enhancer deletion in freshwater sticklebacks led to the loss of pelvic spine armor (Chan et al. 2010). In the hominoid lineage, a transposable element insertion into the intron of a key tail development gene led to an alternative splicing isoform that played a significant role in the loss of tails in humans and apes (Xia et al. 2024). Within *Drosophila*, chromosome-scale cosmopolitan inversions vary in frequency across latitudinal and altitudinal clines, and they have been linked to both thermal adaptation and seasonal adaptation (Kapun and Flatt 2019; Kapun et al. 2023; Nunez et al. 2024).

Despite their outsized role in genomic variation, structural variants have been difficult to study. Their size and location in often-repetitive regions of the genome has made them hard to map and detect using traditional short-read sequencing data. The evolution of long-read sequencing technologies over the last decade has greatly improved our ability to find and analyze structural variants. Long reads are able to capture large structural variants and the flanking genomic regions in single reads, making them detectable and alignable. The addition of extremely long reads has been a key innovation for gapless genome assemblies (Jain et al. 2018; Ebert et al. 2021; Nurk et al. 2022). Furthermore, continuous improvements to sequencing chemistries have increased the accuracy of long-reads to the point where very small variants can be confidently detected (Kolmogorov et al. 2023).

Structural variants are primarily detected through aligning sequencing reads or whole genome assemblies against a reference assembly (Ahsan et al. 2023). Computational methods then identify structural variants by unique mapping signatures within these alignments. Most structural variant calling tools use alignments of long reads to a reference genome. However within the last few years, the process of de novo genome assembly has become more computationally tractable. Furthermore, this new era of haplotype-resolved (or even telomere-to-telomere) assemblies provides a new, highly accurate data type from which structural variants can be called (Ebert et al. 2021). Using these high-quality assemblies, assembly-based methods can detect much larger variants (such as major insertions or inversions) more easily than read-based methods.

Within the last few years, the technical and financial costs of long-read sequencing have decreased such that population-level long-read datasets are feasible to generate for individual labs (De Coster et al. 2021). These datasets are essential for uncovering structural variantion polymorphisms, which better capture the dynamic evolutionary processes that shape structural variant diversity. When generating variant data at the population level, accuracy is paramount to avoid compounding effects of false calls when combining data across many individuals (Ahsan et al. 2023). Though SNP joint genotyping is a wellestablished procedure (Poplin et al. 2018), structural variants can be hard to compare across individuals and require more complex validation strategies (Jeffares et al. 2017; Kirsche et al. 2023; Zheng et al. 2024). While benchmarked structural variant sets have been established for humans (Collins et al. 2020; Zook et al. 2020; Olson et al. 2023), highly validated structural variant callsets do not exist for most other species. Furthermore, many structural variant-calling methods are benchmarked on human data, leaving uncertainty in their accuracy for non-human species. Validation of individual, computationally derived variant calls can be achieved by manually inspecting reads or genome assemblies and their alignments (Belyeu et al. 2018, 2021). While manual validation and curation can identify structural variants with extremely high accuracy, it requires significant effort for genome-scale studies across many individuals (Bertolotti et al. 2020).

A primary determinant of variant-calling accuracy is read length, as exemplified by the structural variation boom that has occurred since moving from short-read sequencing to long-read sequencing. Of the two primary long-read technologies, Oxford Nanopore Technologies (ONT) long-read sequencing can have a much wider read-length distribution than PacBio’s high-fidelity (HiFi) sequencing. While HiFi reads are generally constrained to ∼15kb in length, ONT long reads have no hard limit, and reads larger than 1Mb have been sequenced (Payne et al. 2019). Individual nanopore long-reads within the same sequencing run can range 1000× from hundreds of bases to megabases in length. ONT long-read length is a function of many parameters, including input sample quality, DNA extraction method, and library prep kit (Wang et al. 2021). Given the many factors that can impact long-read length, it is surprising that many structural variant studies do not provide any specific justifications for their long-read lengths beyond technical limitations (Long et al. 2018; Hämälä et al. 2021; Li et al. 2023; Mérel et al. 2023; Shi et al. 2023; Liang et al. 2024). Just as a drastic impact on structural variant calling was seen moving from short reads to long reads, it is likely that the accuracy of structural variant calling significantly varies along the spectrum of read lengths produced by long-read sequencing methods.

In this study, we used the model system of *Drosophila melanogaster* inbred lines to test the impact of different read lengths on the accuracy of automated structural variant calling at the population level. *D. melanogaster* has a rich suite of literature describing functionally adaptive structural variants, including the cosmopolitan inversions (Kennington et al. 2007; Kapun et al. 2016; Kapun and Flatt 2019) and transposable elements (Gonźalez et al. 2008, 2010; Casacuberta and Gonźalez 2013). Transposable elements can vary in length and population frequency, and can be highly similar across copies in the genome (Mérel et al. 2020). Due to these characteristics, transposable element insertions and deletions are hard to align with sequencing data and are challenging to call accurately. Still, transposable elements are estimated to account for a significant proportion of structural variation in *D. melanogaster* (Rech et al. 2022), and thus correctly calling them is a crucial task.

We additionally reasoned that calling structural variants in the compact and gene-dense euchromatic regions of the *D. melanogaster* genome would be an easier task than in larger, intron-heavy and intergenic-region-heavy genomes of vertebrates. ONT long-read sequencing protocols have been developed for *D. melanogaster* to cheaply and efficiently sequence very long (*>*100kb) reads (Kim et al. 2021). *D. melanogaster* has a small (145Mb) genome that makes it financially and computationally tractable for deep long-read sequencing and genome assembly of multiple individuals. Finally, inbred lines are treated as effectively homozygous (i.e. haploid), simplifying many processes from genome assembly to read alignment to variant calling.

Here, we sequenced eight *D. melanogaster* inbred strains to extremely high coverage with ONT using protocols to maximize read length. We then downsampled these deep read pools to generate 30×-coverage read pools with distinct read-length distributions, from which we additionally assembled genomes. We employed five, well-known, automated structural variant callers on each of these distinct read sets and genome assemblies (Figure 1). By manually validating more than 2,300 putative structural variant calls and 16,000 genomic loci across the eight inbred lines, we determined the overall variant-calling accuracy for each read-length distribution. We found that structural variants called by the longest reads and their resulting assemblies were the most accurate, regardless of variant size. For variants smaller (<10kb) than the majority of our long reads, we found that the rest of our read-length distributions were reasonably accurate. Larger variants (≥10kb) could only be accurately called with the ultra-long data. We found that many of the false positives stemmed from misalignments of mobile elements or misalignments in highly complex, repetitive regions.

**Fig. 1.**
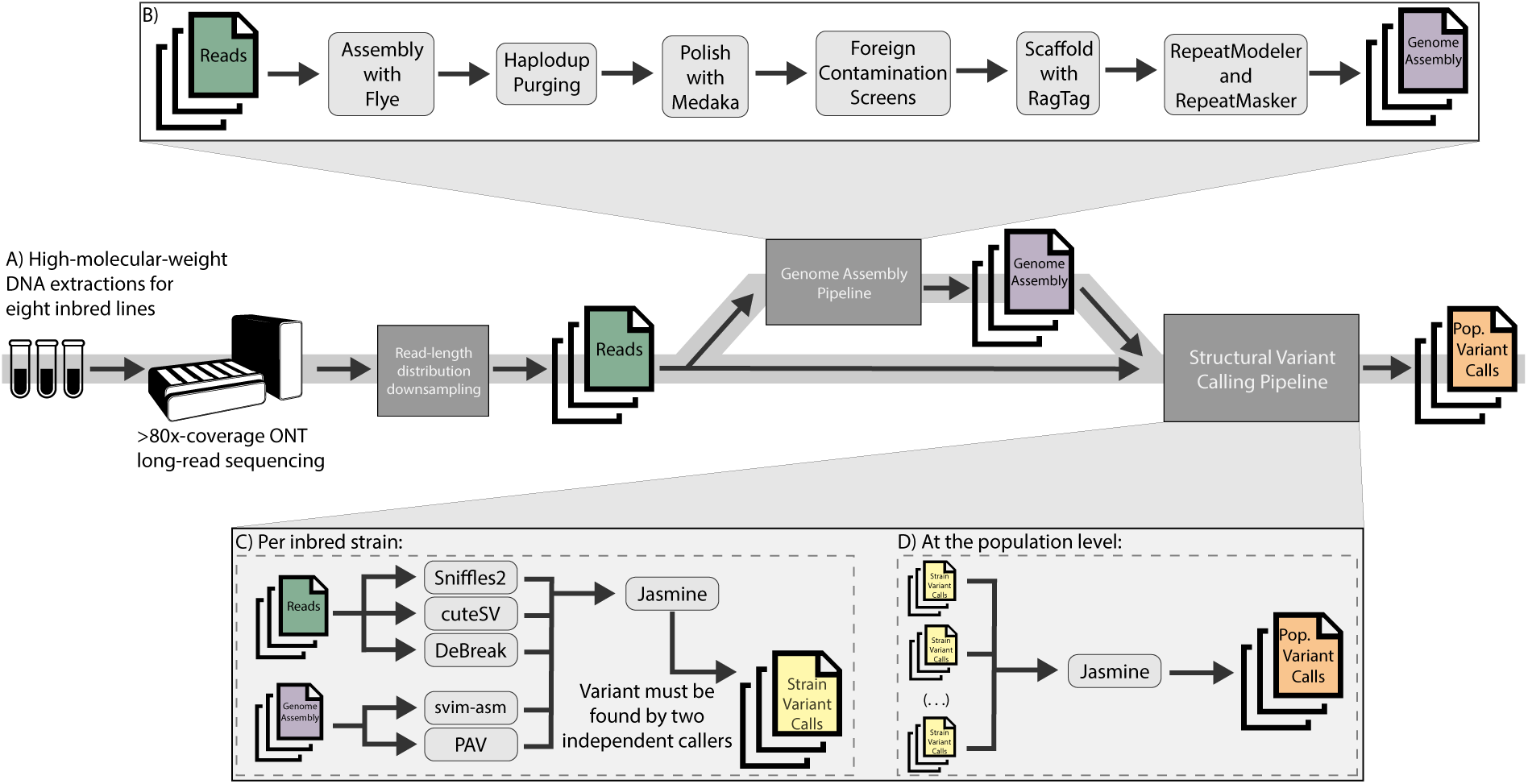
Overview of the methodological workflow. A) Each strain was sequenced to extremely high coverage with ONT long-read sequencing. Various read-length distribution read pools were made by computational downsampling. B) Reads were additionally assembled into genomes. C) Structural variants were called from both read and assembly alignments for each strain and each read-length distribution using five different callers. A variant had to be called by two different callers to be included in the final strain variant set. D) Variant calls from all strains were then merged into a final population variant set. Random variants from the final population set were manually validated.

## Results

### A ladder of read-length distributions

To investigate the effects of long-read length on automated structural variant calling, we performed ONT long-read sequencing on eight inbred *D. melanogaster* lines to generate “master” read pools of ∼80-365x coverage. We then computationally downsampled these reads to generate 30×-coverage pools of different read lengths (Figure 2a). We quantified the read-length distributions using the read N50 statistic, which represents the minimum length of reads such that 50% of the total data are represented in reads that length or longer. We uniformly downsampled the master pools of each line to generate a “standard” readlength distribution, resulting in read N50s ranging from 10kb-19kb (mean=15kb; Table 1). This read N50 range was in line with previous *D. melanogaster* structural variant studies (Chakraborty et al. 2018, 2019; Rech et al. 2022). In this study, we followed the ONT documentation and defined the “ultra-long” distributions as having read N50s greater than 50kb. To generate the ultra-long read-length distributions (read N50s 57kb-87kb; mean = 71kb; Table 1), we enforced a minimum read-length cutoff (35kb-60kb; mean = 52kb) such that we had 30×-coverage of the longest reads from the master read pools. We finally generated a “minimum-10kb” read-length set (read N50s 22kb-38kb; mean = 27kb; Table 1), where all reads less than 10kb were discarded from the master read pools before we performed uniform downsampling to 30×-coverage. The minimum-10kb read pools were designed to mirror the standard read pools but without the shortest reads. This was to control for the fact that the ultra-long read pools had a minimum length threshold, and the presence of shorter reads could create alignment errors that affected structural variant calling more significantly than the gains from increased read lengths.

**Fig. 2.**
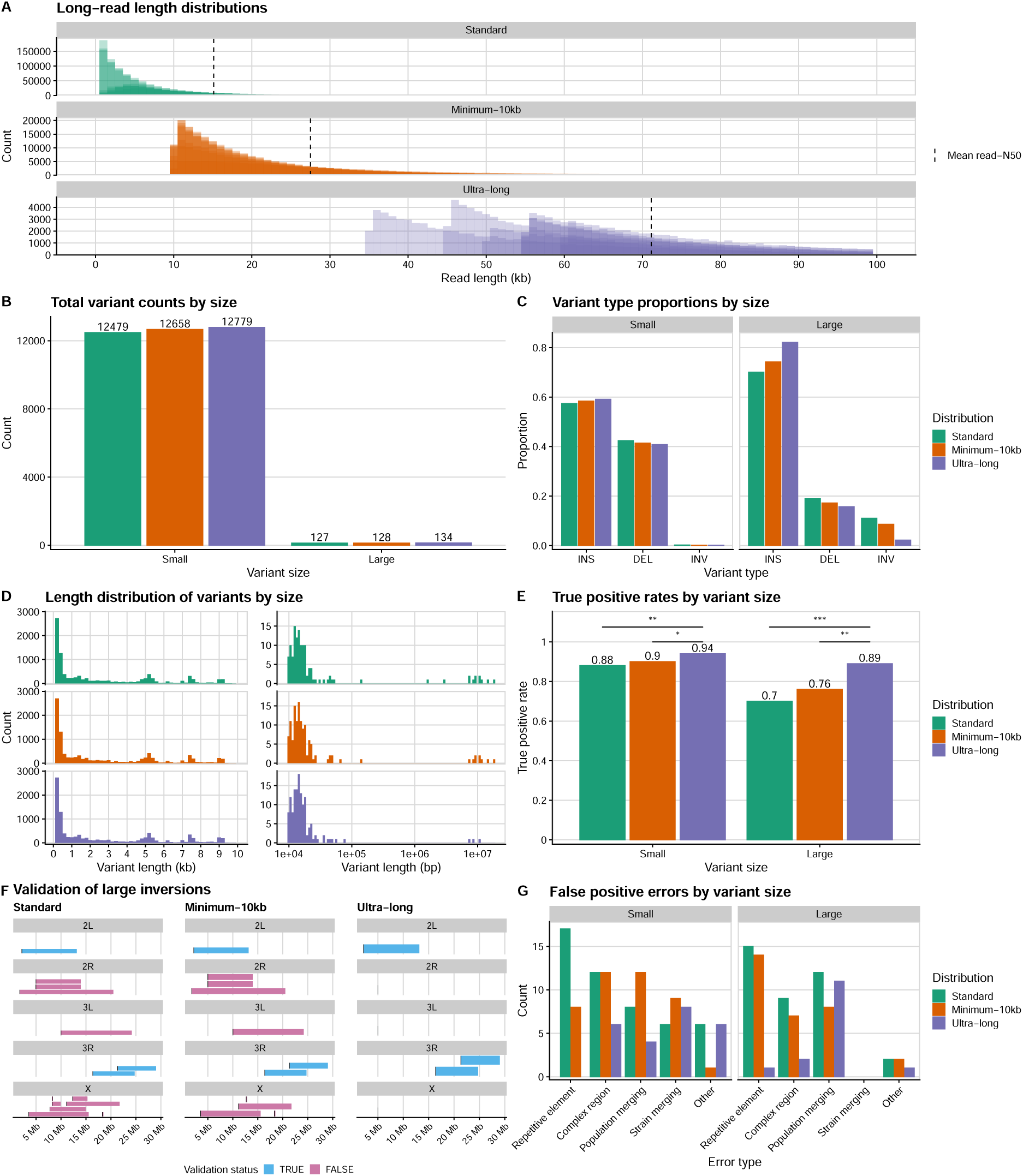
Ultra-long data finds the most comprehensive set of structural variants. A) Read-length histograms of the three, main long-read-length distributions used in this study. Each histogram shows the individual read-length distributions for each of the eight inbred lines. The dashed line represents the mean read N50 of the eight lines for that distribution. For clarity, the histogram only shows reads up to 100kb, however read lengths extend up to 1Mb at minimal frequency. B) The total structural variant counts found by each read-length distribution, broken down by variant size. Small variants were defined as being less than 10kb in length, while large variants were equal to or longer than 10kb. C) The relative proportions of variant types found by each read-length distribution, broken down by variant size. D) The variant length histograms for small (left) and large (right) variants found by each read-length distribution. E) The true positive rate of each read-length distribution as determined by manual validation, broken down by variant size. F) Chromosomal maps of all large inversions found by each read-length distribution, colored by whether those inversions are real, as determined by manual validation. G) The false positive errors as determined by manual validation for each read-length distribution, broken down by variant size. Repetitive element errors stemmed from incorrect alignment of mobile or repeat elements. Complex region errors were caused by incorrect alignments in highly repetitive regions or in regions with intersecting structural variants. Population merging errors were caused by incorrect genotyping of specific variants across the eight strains. Strain merging errors were when very small indels within a strain were incorrectly merged together and called as a larger variant. All other errors were binned into the “Other” category. * = p *<*.05; ** = p =.001; *** = p *<*.0001

**Table 1.** Sequencing and assembly statistics for each of the read-length distributions averaged across the eight inbred lines. Note that the master read pools and the short-read pools were not assembled. For line-specific statistics, please see Supplementary Table 1.

| Distribution | Coverage<br>(145Mb) | Minimum read<br>length (bp) | Read<br>N50 (bp) | Contig<br>N50 (Mb) | Total scaffold<br>length (Mb) | Scaffold<br>N50 (Mb) | Total<br>scaffolds | Busco<br>completeness |
| --- | --- | --- | --- | --- | --- | --- | --- | --- |
| Master | 237.8 | - | - | - | - | - | - | - |
| Standard | 31.6 | 5 | 15085 | 13.6 | 129.6 | 23.8 | 43.5 | 99.5 |
| Minimum-10kb | 29.7 | 10000 | 27448 | 14.6 | 132.3 | 24.4 | 19.5 | 99.5 |
| Ultra-long | 30.3 | 51875 | 71184 | 17.8 | 137.1 | 25.1 | 13.5 | 99.5 |
| Ultra-long-20× | 20.2 | 51875 | 71206 | 19.3 | 134.7 | 24.7 | 14.8 | 99.5 |
| Ultra-long-10× | 9.9 | 51875 | 71254 | 9.8 | 133.6 | 24.7 | 15.1 | 99.3 |
| Short-read | 46.6 | 10 | 150 | - | - | - | - | - |

### High-quality genome assemblies from long-read data

To make use of assembly-based structural variant callers, we assembled each of the standard, ultralong, and minimum-10kb read-length pools into full genomes (see Materials and Methods). Overall, these assemblies were highly contiguous (Table 1), with the standard assemblies having scaffold-N50s between 23.2Mb-25Mb. The ultra-long reads generated near-chromosome-level assemblies, with scaffold-N50s between 24.3Mb-26.1Mb. The minimum-10kb assemblies fell in the middle with scaffold-N50s between 23.8Mb-25.5mb. High contiguity, especially in euchromatic regions, resulted in very high degrees of benchmarking universal single-copy ortholog (BUSCO) completeness (all *>*99%).

### Ultra-long reads and assemblies greatly increase accuracy for structural variant calling

To call structural variants from long-read data, we employed five, well-known, automated structural variant callers. Sniffles2 (Smolka et al. 2024) and cuteSV (Jiang et al. 2020) use strictly read-based alignment approaches, DeBreak (Chen et al. 2023) uses a read-based approach coupled with local reassembly around the putative structural variant, and svim-asm (Heller and Vingron 2021) and PAV (Ebert et al. 2021) utilize assembly-based alignment approaches. For each read-length distribution, we aligned the 30×-coverage read pools and the resulting assemblies to the *D. melanogaster* reference genome (v6.58). We limited our analyses to insertions, deletions, and inversions, as these variant types were universally recognized by our variant callers. Most of the callers we used also had the ability to detect duplications, and we converted those calls to insertions so that they could be properly merged.

All of our alignments and structural variant calls were subject to multiple filtering steps, including minimum alignment scores, minimum and maximum genome coverage, and “PASS” and “PRECISE” variant flags (see Materials and Methods). Leveraging a strategy employed by previous studies to reduce the false positive rate, we required all variants to be independently found by at least two automated callers (Li et al. 2023; Mérel et al. 2023; Couper et al. 2025; Jensen et al. 2025). In line with other *D. melanogaster* structural variant studies (Chakraborty et al. 2018, 2019; Rech et al. 2022), we chose to only look at structural variants found in the euchromatic regions of the genome (Supplementary Table 2). We chose conservative filtering steps throughout our variant calling pipeline as false negative calls at the strain level could lead to significant artifacts at the population level when variant calls were merged.

As no benchmark structural variation datasets currently exist for *D. melanogaster*, we validated structural variant calls using a manual approach where we visualized the ultra-long read alignments of each strain at the putative variant locus with Jbrowse2 (Diesh et al. 2023). A variant was considered a true positive call if it was manually found to be exclusively present in the same inbred lines as found by the callers. We divided our manual validation into two categories, looking at the performances of the callers on “small” structural variants, which we defined as variants with a length less than 10kb, and on “large” structural variants greater than or equal to 10kb. We chose this threshold as it was just smaller than the read N50s of our shortest long-read distribution and because the largest transposable elements in *D. melanogaster* are just under 10kb in length.

When looking at small structural variants, we found subtle differences in the types of variants detected by each of the three read-length distributions. The standard data found 300 (2%) fewer structural variants than the ultra-long data (Figure 2b). We next checked the proportions of our variant types within these counts, and we found that the ultra-long data called the largest proportion of insertions and the smallest proportion of deletions, while the standard data showed the inverse (Figure 2c). Though the small inversion proportions were less than one percent for all read-lengths, the standard data found the most (19) while the ultra-long data found the fewest (14). Finally, the variant length histograms showed no qualitative differences between the datasets (Figure 2d). Each histogram showed that most structural variants were less than 1000bp. There were additional modes representing variation caused by different mobile element families (Mérel et al. 2020), a pattern that has been observed in other species (Weissensteiner et al. 2020; Ebert et al. 2021; Li et al. 2023).

Despite the subtle differences in the makeup of variants called by the different distributions, the smallvariant true-positive rates for the standard and ultra-long read-length distributions differed by nearly 10% (Figure 2e). We randomly sampled and validated 400 small variants from each of the callsets. Of the 400 standard, small structural variants, 350 (87.5%) were true positive calls. The ultra-long data found a statistically significant increase in true positive calls (chi-square test; p=.001 against standard), with 376 (94%) of the 400 small variants being true positives. While the standard read-length distribution accurately called many structural variants less than 10kb in length, the ultra-long data ultimately captured more small variants and with higher accuracy.

We repeated our analyses, next focusing on large structural variants with lengths of at least 10kb. Again, each of the read-length distributions recovered very similar numbers of total large variants, with the standard data finding just seven fewer (5%) than the ultra-long data (Figure 2b). The proportion of insertions differed, with 110 (82%) of the ultra-long variants being insertions, while the standard data found only 89 (70%) insertions (Figure 2c). The standard large variants included 14 (11%) inversions, while the ultra-long data found just three (2%) inversions. The standard data was also composed of more deletions (24; 19%) than the ultra-long data (21; 16%). The large-variant length histograms showed almost all variants found by the ultra-long data were in the 10-100kb range, while the standard variants had a second mode around 10Mb (Figure 2d).

Ultimately, the large-variant true-positive rate was much higher for the ultra-long data than for the standard dataset. We manually validated all of the large structural variant calls from the standard and ultra-long data. 121/134 (90%) of the ultra-long large variants were found to be real (Figure 2e). Strikingly, the large variant calls from the standard data were much less accurate (chi-square test; p *<*.0001) as only 89/127 (70%) were true positives. Again, the ultra-long data returned the largest and most accurate set of structural variants. We specifically compared the sets of large structural variants called by the two datasets and found that about one quarter of variants were unique to the ultra-long data (35/134) and to the standard data (30/127). 30 of the 35 (86%) of the unique ultra-long variants were validated as real, while just 5 of the 30 (17%) unique standard variants were true positives. We found that the true positive calls unique to the standard dataset occurred in genomic regions where the ultra-long data had abnormally low coverage.

While the ultra-long data contained reads that were on average ∼7x longer than the standard reads, the ultra-long data also contained no reads less than 35kb. It was possible that the improvements seen with the ultra-long data came from the lack of small reads, which would often align poorly, rather than an excess of longer reads. We repeated our analyses with the minimum-10kb data, expecting the minimum-10kb data to show similar results to the ultra-long data if indeed that was the case. We found that the minimum-10kb results always fell between the standard and ultra-long values (Figure 2). Ultimately, the minimum-10kb true positive rates for both small (358/400; 89.5%) and large variants (97/128; 76%) were significantly lower than the ultra-long true positive rates (chi-square test; p < 05 for small; p = .001 for large), suggesting that the removal of the shortest reads was not the most significant factor to the ultra-long data’s high accuracy. Rather, the length of the reads in the ultra-long dataset was crucial to correctly aligning and assembling problematic regions with structural variants.

Chromosomal inversions are well-characterized in *D. melanogaster* as being adaptive variants, thus it is of particular importance to be able to call them correctly. Both the standard (14) and minimum-10kb (11) called significantly more large inversions than the ultra-long data (3) found. Strikingly, all three of the inversions found in the ultra-long data were known cosmopolitan inversions, and our manual validation confirmed their presence in our inbred lines. While the other two read-length distributions also found these three cosmopolitan inversions, none of the remaining inversion calls were real (Figure 2f).

### Misalignments of transposable elements and complex repetitive regions primarily cause false positives in long reads

To better understand why automated structural variant callers make errors, we categorized the 199 false variant calls (both small and large) from our three long-read datasets into five categories (Figure 2g). In more than half of the false positives, reads containing mobile or repeat elements (“repetitive element” errors) or reads in complex regions were incorrectly aligned, leading to false structural variant alignment signatures. Complex regions were defined as loci with non-repeat-element repetitive sequence (such as the Stellate gene cluster (Tulin et al. 1997)) or regions with multiple structural variants intersecting each other (such as the Cyp6g1/2 locus (Schmidt et al. 2010)) (Supplementary Figure 1). These types of errors stemmed from having reads too short to fully span both the repetitive or complex loci and their surrounding genomic regions. As a result, the aligner incorrectly mapped or split-mapped reads. Repetitive element errors led to all false, large inversion calls. At each of the fake breakpoints of the false inversions, there were either transposable element insertions or deletions. Reads that could not span the full transposable element event at one “breakpoint” were incorrectly split-mapped to the event at the other “breakpoint”, creating a fake alignment signature that looked like an inversion to the variant callers (Supplementary Figure 2). The ultra-long dataset accounted for just 8% (9/106) of false positives stemming from repetitive elements or complex regions, with eight of those nine calls being complex region errors. The single ultra-long “repetitive element” error was from an incorrectly mapped sequence ∼80kb in length that was almost exclusively transposable elements (Supplementary Figure 3).

The secondary source of false positive variant calls was merging errors, either at the population level or strain level. “Population merging” errors occurred when dissimilar variants at the same genomic locus across different strains were merged and considered the same variant across those strains (Supplementary Figure 4). 31 of the 55 “population merging” errors occurred in large variants, and 73% (11/15) of the ultra-long large false positives were due to this type of error. A number of these cases involved very large structural variants (*>*20kb) in regions with variable read coverage across the strains, leading to these variants being correctly called in only a subset of the strains. “Strain merging” errors occurred when very small indels (often < 50bp) in one strain were incorrectly grouped together to form a single, larger variant call (Supplementary Figure 5). Automated callers will often left-align or right-align variants in repetitive regions, however in non-repetitive regions this alignment shift can incorrectly merge small variants together. We found all 23 of the “strain merging” errors when validating “small variants”. None of the read-length distributions were significantly biased for or against either of the types of merging errors.

We binned the rest of the false positive error calls into “other” errors, which constituted less than 10% of all false positive calls. These false positives were due to errors from sequencing, from genome assembly, from genomic regions with low read coverage, and from small, stochastic bumps or dips in read coverage. For example, we observed a large, inverted duplication call perfectly captured in a single read. In reality, this could have been due to the “unzipped”, second strand of the DNA molecule being immediately sequenced in the same nanopore after the first strand and then incorrectly counted as a single molecule of DNA. In a limited number of cases (< 2 %), we could not determine the cause of the false positive variant call.

### Long-read structural variant calling loses accuracy at lower coverage

Generating ultra-long reads from *D. melanogaster* is a relatively time-consuming and expensive process as compared to other long-read protocols. To see if we could make ultra-long reads more accessible, we uniformly downsampled our ultra-long read pools from 30×-coverage to 20×-(ultra-long-20×) and 10×-coverage (ultra-long-10×) and then assembled genomes for each of the eight inbred lines (Table 1). We then called structural variants from the ultra-long-20× and ultra-long-10× data using the same pipeline and compared them to our original ultra-long variant calls.

The 20×-coverage data called just a couple hundred (2%) fewer total variants than the 30×-coverage ultra-long data, while the 10×-coverage data called almost 10% (1,150) fewer structural variants (Figure 3a). Despite the spread in total variants called, all three ultra-long distributions had very similar proportions of variant types and showed very similar variant length distributions (Figure 3b). Each of the different coverages successfully recovered the secondary modes corresponding to different mobile element families. Like the ultra-long dataset, both the 20×-coverage and 10×-coverage datasets called just a handful of structural variants larger than 1Mb (Figure 3c). In spite of these broad similarities, the lower-coverage datasets did not match the accuracy of the 30×-coverage ultra-long dataset. We manually inspected all large variant calls for the ultra-long-20× and ultra-long-10× datasets and found that the ultra-long-20× variant set had an 82% true positive rate, while the ultra-long-10× set had just a 72% true positive rate, which were both lower than the 30×-coverage dataset’s accuracy of 89% (Figure 3d). Only the ultra-long-10× data’s true positive rate (chi-square test; p = .001) was statistically significantly lower than that of the original 30×-coverage’s rate, however neither of the lower coverage datasets was able to recover all of the large inversions found by the original 30×-coverage data (Fig. 3e).

**Fig. 3.**
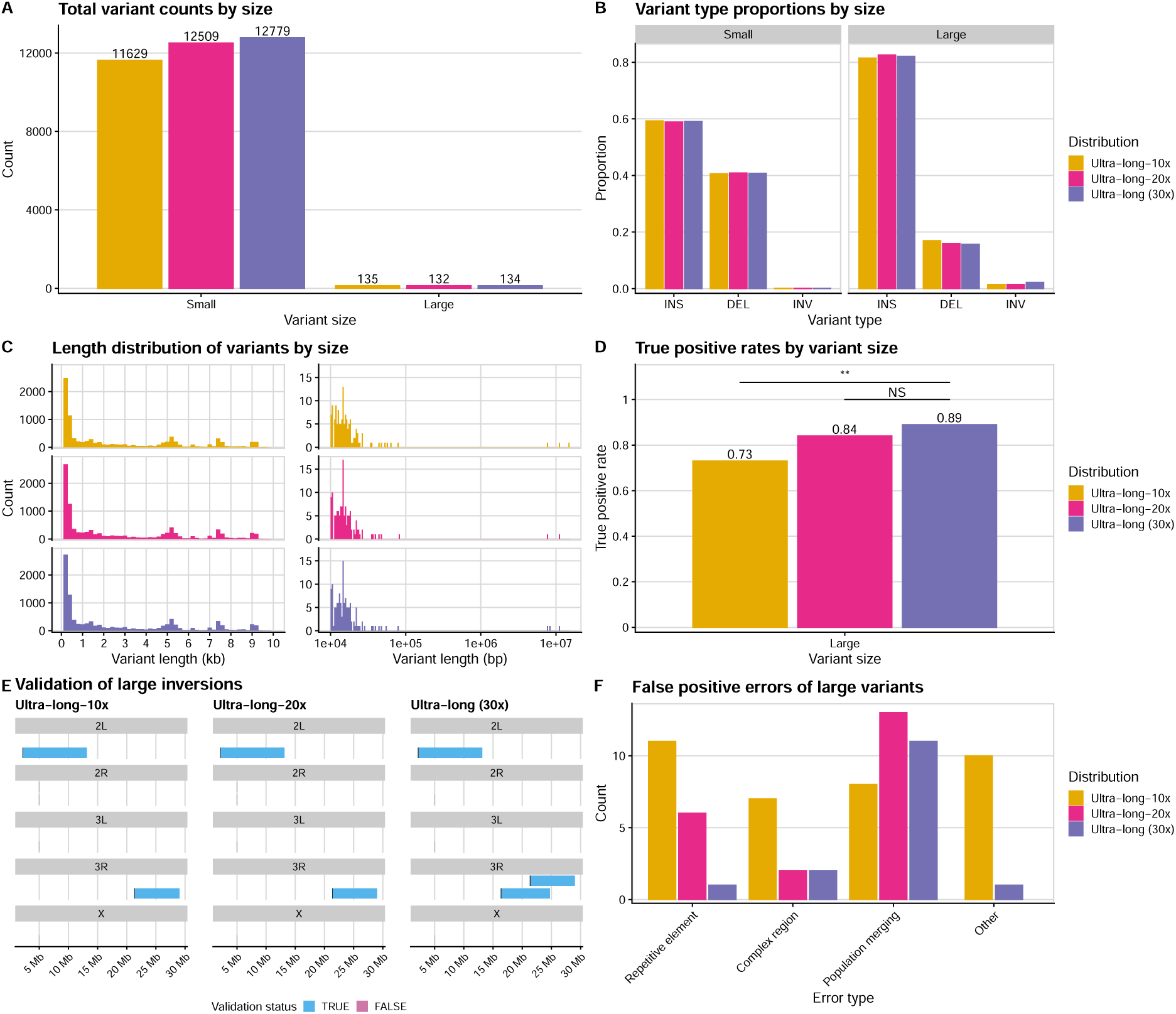
Ultra-long data loses variant-calling accuracy as coverage decreases. A) The total structural variant counts found by each ultra-long-read coverage distribution, broken down by variant size. B) The relative proportions of variant types found by each coverage distribution, broken down by variant size. Small variants were defined as being less than 10kb in length, while large variants were equal to or longer than 10kb. C) The variant length histograms for small (left) and large (right) variants found by each coverage distribution. D) The true positive rate for large structural variants of each coverage distribution as determined by manual validation. E) The false positive errors of large variants as determined by manual validation for each ultra-long coverage distribution. Repetitive element errors stemmed from incorrect alignment of mobile or repeat elements. Complex region errors were caused by incorrect alignments in highly repetitive regions or in regions with intersecting structural variants. Population merging errors were caused by incorrect genotyping of specific variants across the eight strains. Strain merging errors were when very small indels within a strain were incorrectly merged together and called as a larger variant. All other errors were binned into the “Other” category. NS = Not Significant; ** = p = .001

A substantial fraction of these errors, especially for the 10x-coverage data, came from misalignments from repetitive elements and complex regions (Figure 3f). The decreases in coverage meant the aligner had to work with fewer reads that spanned specific loci. Additionally, with an average of just 10 reads per genomic locus, the relative frequency of poor alignments was much higher, leading to more errors. We observed an increase in the number of “population merging” errors in the ultra-long-20x data. In these cases, the coverage of the variant was unevenly decreased across different strains, leading to some strains receiving incorrect calls. Finally, 10 false calls from the ultra-long-10x data were categorized in “other” errors. Many of these were seemingly heterozygous variants that had too low coverage in the 10x-coverage dataset to be accurately called. Even though we were using inbred lines, it is well known that inbred lines can maintain heterozygosity of both SNPs and structural variants (Frankham et al. 1993; Rumball et al. 1994; Powell and Evans 2017; De Kort et al. 2022).

### Short-read data do not accurately capture the full breadth of structural variation

The vast majority of existing genomic data is short-read data, so we additionally tested the capabilities of short-read data for accurately calling structural variants. We sequenced each of the eight lines with Illumina to generate standard, 2 x 150bp paired-end, short-read data to an average of 46x-coverage (Table 1). For the short-read data, we called variants with four established, short-read-based callers, Manta (Chen et al. 2016), Delly2 (Rausch et al. 2012), Lumpy (Layer et al. 2014), and GRIDSS2 (Cameron et al. 2017). Again, we merged with Jasmine, and we required all variants to be independently found by at least two of the four automated callers. We compared to the resulting structural variant calls to those found by the standard and ultra-long long-read datasets.

Immediately, we found differences between the short-read data and the long-read datasets, as the short-read data called just 4,697 small variants, or 37% of what the long-read datasets found (Figure 4a). When looking at the proportions of small variant types found, the short-read data had a striking lack of insertions (14%) and a huge percentage of deletions (84%) (Figure 4b). The length histogram of the shortread variants did not display secondary modes corresponding to mobile element families like the long-read distributions had shown (Figure 4c). We manually validated 400 random, small structural variants from the short-read data and found that the accuracy (81%) was worse than all of the long-read callsets (chi-square test; p *<* 1e-4 against ultra-long data) (Figure 4d). With both a lower true positive rate and a severe deletion bias, structural variant calling with short-read data failed to create an accurate picture of the small structural variant landscape.

**Fig. 4.**
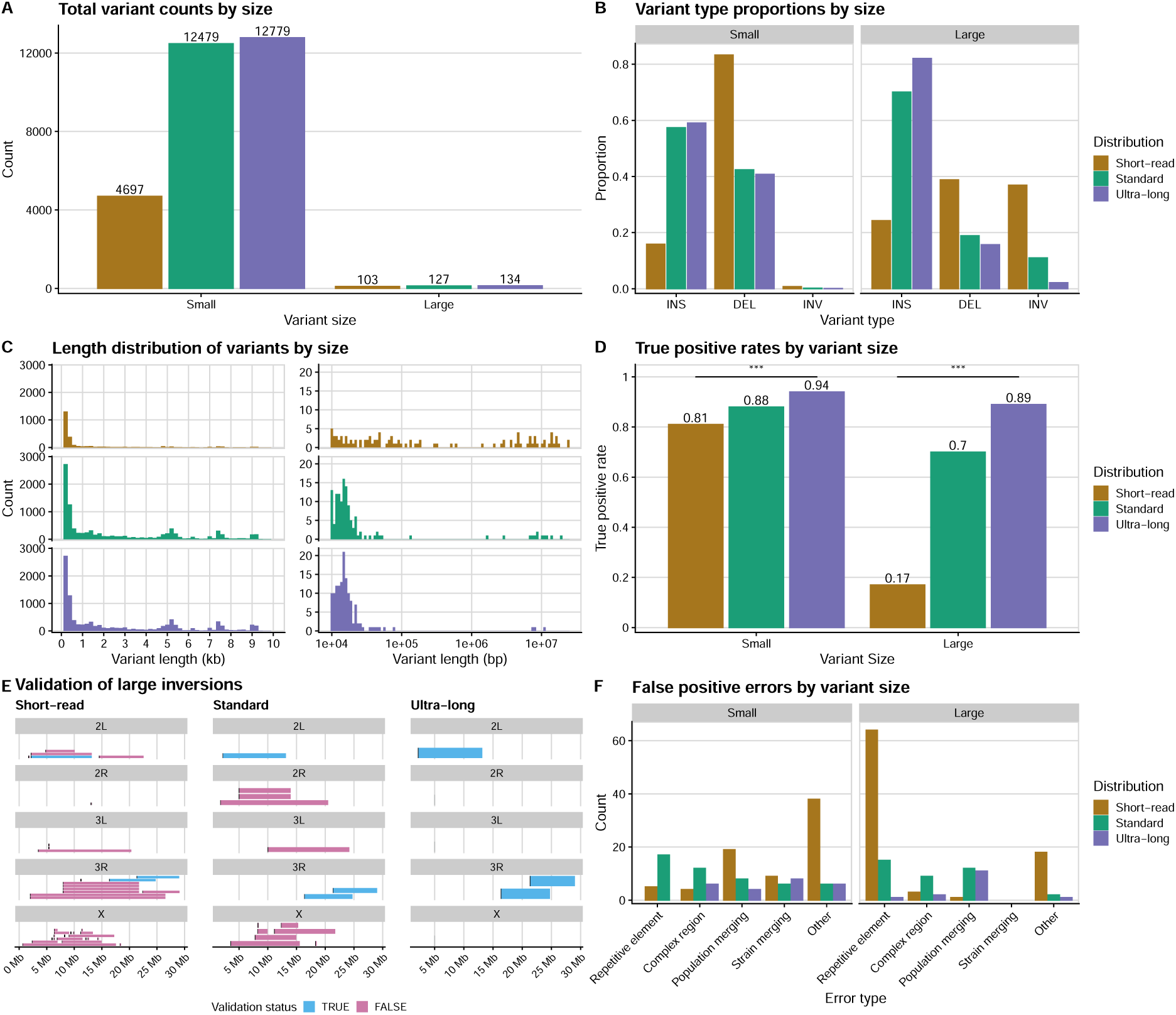
Short-read data recovers a biased set of variants at low accuracy. A) The total structural variant counts found by each read-length distribution, broken down by variant size. B) The relative proportions of variant types found by each read-length distribution, broken down by variant size. Small variants were defined as being less than 10kb in length, while large variants were equal to or longer than 10kb. C) The variant length histograms for small (left) and large (right) variants found by each read-length distribution. D) The true positive rate for large structural variants of each read-length distribution as determined by manual validation. E) The false positive errors of variants as determined by manual validation for each read-length distribution, broken down by variant size. Repetitive element errors stemmed from incorrect alignment of mobile or repeat elements. Complex region errors were caused by incorrect alignments in highly repetitive regions or in regions with intersecting structural variants. Population merging errors were caused by incorrect genotyping of specific variants across the eight strains. Strain merging errors were when very small indels within a strain were incorrectly merged together and called as a larger variant. All other errors were binned into the “Other” category. *** = p *<*1e-4

When comparing the large structural variant calls, we again found that the short-read data severely under-called insertions and over-called deletions and inversions compared to the long-read data, even though the overall numbers were much closer (Figure 4b). The short-read data notably called the most large inversion (38) out of any read-length distribution. The large-variant size distribution from the shortread data appeared markedly different from any of the long-read distributions, with a much more uniform distribution of variants up to, and past, 10Mb (Figure 4c). We manually validated each of the 103 large, short-read variants and found that just 17 (17%; chi square test; p *<* 1e-4 against ultra-long data) were real variants (Figure 4d). Nearly all (35/38) of the large inversions called by the short-read data were false positives (Figure 4e). The vast majority of the large false positive calls came from spurious signatures generated by misaligned TE and DNA repeat sequences (Figure 4f). The short-read variant calls had a higher incidence of “other” errors as compared to the long-read data, especially for small variants. We found these to be largely the same types of errors, as found in the long-read data, just at a higher frequency. There did not appear to be an error type unique to short-read data.

## Discussion

We report significant shifts in automated structural variant calling accuracy at the population level when systematically varying read length within *D. melanogaster*. Our longest long-read-length distribution, the ultra-long reads (minimum read-length = 35kb; average read N50 = 71kb), called more structural variants, and at a higher accuracy, than any other read-length distribution. We found that the total number of called structural variants, the number of called insertions, and the true-positive rate all increased as the read N50 increased. While all long-read datasets called structural variants less than 10kb in length with at least 88% accuracy, only the ultra-long dataset maintained similar accuracy with variants longer than 10kb.

In this study, we manually validated more than 2,300 putative structural variant calls at more than 18,000 genomic loci across eight inbred strains. Our manual validation not only allowed us to assess the accuracy of computational variant calls, but also let us determine the causes behind the false positives. In all datasets except for the ultra-long distribution, misalignments from repetitive elements and complex genomic regions caused more than half of all false positive calls. In these cases, the reads were not long enough to both fully span the problematic region and capture enough surrounding genomic context. As a result, the aligner incorrectly mapped repetitive elements, leading to false structural variant signatures for the callers.

In *D. melanogaster*, the longest mobile elements approach 10kb, and our results suggest that reads significantly longer than the largest repetitive elements are required to accurately map them. Our ultralong reads, with a minimum read-length of 35kb, were able to map these elements, suggesting that reads at least 3x longer than the largest repetitive element are required to avoid variant-calling errors from these elements. Additionally, only the 30x-coverage of ultra-long reads was able to accurately call the three cosmopolitan inversions. The two low-coverage ultra-long datasets each recovered just two out of the three inversions. These results further highlight that accurate structural variant calling is a species-specific problem, as the repeat element landscape is highly variable across taxa. *D. melanogaster* harbors a relatively small number of mobile elements, however different copies of some transposable elements can have very high sequence similarity, hindering accurate alignment without additional genomic context.

The primary type of error for the ultra-long reads, and a secondary error contributor to all other readlength datasets, was error from merging variants across individuals. In these cases, structural variant calls either were incorrectly assigned to strains that did not contain the variant or were absent from strains that actually did contain the variant. Merging structural variant calls across different individuals is a complex problem, and our results highlight the importance of developing algorithms that include multiple types of evidence for merging, including breakpoints, variant length, and variant sequence content.

Variant-calling errors can have a significant impact on downstream population genetics analyses. High error rates introduce many false, low-frequency polymorphisms which can potentially mislead inferences of mutation rate and selection. Similarly, variant merging errors at the population-level can skew the site frequency spectrum of structural variants. Variant-calling biases can similarly change allele frequencies and affect inferences of selection for different types and sizes of variants.

The ability to accurately call a much fuller breadth of structural variants, without biases for variant length or type, opens up new possibilities to answer long-standing questions in population genomics and genome evolution. For instance, without variant-type biases, we can accurately compare the selective forces acting on different structural variants. We can capture and assemble multiple copies of gene duplications, including those that are still polymorphic, and accurately delineate the accumulation of substitutions in different copies to infer the selective pressures on newly duplicated genes (Kondrashov 2012). While many examples of adaptive structural variation have been found through concerted efforts on specific genomic loci (Chan et al. 2010; Xia et al. 2024; Dodge et al. 2024), there are many regions across the genome, found by GWAS or other genome-wide statistical scans using SNPs, that are implicated in selective sweeps (Garud et al. 2015), allelic series (Schmidt et al. 2010), and rapid and seasonal adaptation (Rudman et al. 2022; Bitter et al. 2024) which could be underpinned by now-detectable, functionally-relevant structural variation. We firmly believe that accurate, population-level calling of structural genomic events will allow the field to make key discoveries that are currently obscured by the almost-exclusive focus on single nucleotide polymorphisms.

## Methods

### DNA Extraction and long-read sequencing

We sequenced *D. melanogaster* isofemale, inbred lines originally collected from Linvilla Orchard, PA, in 2011 (Behrman et al. 2015) (Supplementary Table 3). Flies for this study were reared at 25C on cornmeal-molasses fly media with a 12-hour day-night light cycle. Female flies were sexed and starved one day prior to extraction.

We followed previously published *D. melanogaster* long-read sequencing protocols from our lab (Kim et al. 2021, 2024). Briefly, 100-400 female flies were homogenized using Kimble Kontes Dounce homogenizers. The homogenate was centrifuged to pellet debris and separate out nuclei. Nuclei were then resuspended in a lysis buffer and incubated at 50C for ∼4 hours, mixing the tubes by gentle inversions every 45 minutes. The DNA was then purified from the lysate using a standard phenol-chloroform extraction. The lysate was mixed with an equal volume of phenol-chloroform-isoamyl alcohol in phase-lock gel tubes, centrifuged for eight minutes, and inverted for eight minutes on a rocker. An equal volume of chloroform was then added to the phase-lock gel tubes, and again the tubes were centrifuged and inverted for eight minutes each. We precipitated the DNA out of solution by adding 10% volume of 3M NaOAc and 2x volume of cold, 100% ethanol to each tube and gently rocking and inverting the tubes to mix. A wide-bore pipette tip was used to transfer the precipitated DNA to new tubes where it was subsequently washed twice with 70% ethanol. The DNA was pelleted and allowed to air dry for a couple of minutes before being resuspended in 60ul of 1 x Tris-EDTA buffer at 50C for one hour and then for a couple of days at 4C. If the DNA was not fully resuspended after a few days, the DNA was sheared by slowly pipetting twice with a P1000 tip and then allowed a few more days to resuspend. DNA concentration was quantified with Qubit, and Nanodrop absorption ratios were checked to ensure that 260/280 was greater than 1.8 and 260/230 was greater than 2.0.

We prepared the nanopore library following the ONT Ligation Sequencing Kit (SQK-LSK114) protocol, with a couple major modifications. First, we started with ∼4.5ug of DNA as opposed to the recommended 1ug of DNA. Second, we used the Circulomics Short Read Eliminator (SRE) buffer before starting the official Sequencing Kit protocol to remove the shortest DNA fragments. We then performed the Sequencing Kit’s DNA Repair and End-prep steps using half-reactions to save reagents, without loss of quality in the final product. We also performed the adapter ligation and clean-up steps using half-reactions. We additionally let the adapter ligation reaction incubate for ∼30 minutes instead of the prescribed 10 minutes. Finally, we performed a second round of short-read elimination using the Circulomics SRE buffer. Critically, we washed the DNA with SFB or LFB (interchangeable) from the Sequencing Kit because ethanol denatures the nanopore motor proteins. We loaded ∼350ng of prepared library onto R10.4.1 flow cells and sequenced on an ONT PromethION 2 following ONT’s protocol with live basecalling set to fast mode. For further discussion and justification of these protocols, please refer to Kim et al. (2021, 2024).

### Short-read sequencing

To short-read sequence each of the inbred lines, we used the same DNA extracted for the long-read library preps. ∼10ng of DNA were prepared with the Illumina DNA library preparation kit. We followed the standard protocol, but we used one-fifth reactions to save reagents at no cost to the final quality. All libraries were amplified with 6 PCR cycles. The prepared libraries were then cleaned with beads and pooled by concentration. We checked the fragment size distributions with BioAnalyzer, and the concentration of the library pools were quantified via qPCR by Admera Health. Sequencing was performed on the NovaSeq X Plus by Admera Health to 30x-60x coverage. One inbred line (dmel19) was sequenced a second time to reach the desired coverage. We used BBtools (B. 2014) for adapter trimming the short-read data.

### Basecalling and read-length distribution generation

We long-read sequenced the eight inbred lines to extremely high depths (80-365x; master pools) so that we would have high-enough coverage of ultra-long reads. We basecalled our reads using Dorado (model: “dna r10.4.1 e8.2 400bps sup@v5.0.0”) and filtered out reads with a quality score less than 10. To generate our different read-length distribution pools for each inbred line, we downsampled the total pools of reads in various ways. We used the program seqtk (https://github.com/lh3/seqtk) for downsampling.

To generate the standard pool of reads, we used “seqtk sample” to uniformly and randomly downsample the master pool to 30×-coverage. To generate the ultra-long set of reads, we used “seqtk seq” with the “-L” argument to set the minimum read length such that we only kept the longest reads that resulted in 30×-coverage. For the minimum-10kb pools, we first used “seqtk seq” with the “-L” argument to remove all reads less than 10kb from the total pools. Then we used “seqtk sample” to uniformly and randomly downsample the remaining reads to 30×-coverage. For the lower-coverage ultra-long distributions, we uniformly downsampled the ultra-long read pools to 20×-coverage and 10×-coverage. We used the program seqkit to find the summary statistics of each of our downsampled read pools.

### Resequencing for proper standard-length long reads

There were three lines (dmel12, dmel19, dmel21) whose sequencing runs generated much higher proportions of long reads. As a result, the random, uniform downsampling used to create the standard read-length distributions consistently generated pools with read N50s 2-3x higher for these lines. As consistent read-lengths were paramount to our study, we elected to resequence these three lines using the standard long-read protocol from Kim et al. (2021). All other read-length distributions for these three lines were computationally generated from the master pools like the other lines.

### Long-read assembly pipeline

We followed the haploid assembly pipeline described by Kim et al. (2024). We fully assembled genomes for each of the long-read distributions of each inbred line. Briefly, basecalled reads were assembled with Flye (Kolmogorov et al. 2019) using the “–nano-hq –read-error 0.03” parameters. We removed haplotigs from the contigs using purge dups (Guan et al. 2020), and we then polished with Medaka for one round. Contaminant sequences and adapters were found and removed using the NCBI Foreign Contamination Screen (Astashyn et al. 2024). We scaffolded the cleaned contigs against the *D. melanogaster* reference assembly (v6.58) using RagTag (Alonge et al. 2022). Repetitive sequences were found and soft-masked with RepeatModeler (Smit and Hubley 2008) and RepeatMasker (Smit et al. 2013). Assembly metrics were found using the command “gt stat” from the program GenomeTools (Gremme et al. 2013). We determined BUSCO completeness with the program compleasm (Huang and Li 2023), using the “diptera odb10” lineage.

### Long-read structural variant calling

All long-read read pools and assemblies were aligned to the *D. melanogaster* reference assembly (v6.58) using minimap2 (Li 2018). Reads were mapped with the parameter “-x map-ont” and assemblies were mapped with the parameter “-x asm5”. We chose five well-known, structural variant callers that used a variety of algorithms. Structural variants were identified for each read-length distribution in each strain with sniffles2 (Smolka et al. 2024), cuteSV (Jiang et al. 2020), DeBreak (Chen et al. 2023), svim-asm (Heller and Vingron 2021), and PAV (Ebert et al. 2021). We required a minimum read-mapping quality of 20 and a minimum variant length of 50bp for all callers except PAV, which has its own read-mapping quality thresholds.

For each read-length distribution, structural variants were called in each inbred strain by all five callers, resulting in five independent variant calling format (VCF) files per strain. We used bcftools to filter each of the five VCF files to only keep variants with the “FILTER=PASS” and “PRECISE=1” flags. We additionally filtered out all variants that did not overlap the euchromatic regions as defined in Chakraborty et al. (2018) (Supplementary Table 2). We only kept insertions, deletions, duplications, and inversions. We then used a custom script to recode duplications as insertions in the VCF files. This was done to allow for proper merging, as duplications were identified as duplications by some callers and insertions by others. We merged the five VCF files for one strain into a single VCF file using Jasmine (Kirsche et al. 2023), and we filtered out all structural variants that were not called by at least two independent callers (Jasmine parameter “min support=2”). Finally, for each read-length distribution we merged the eight strain VCF files into a single population VCF file using Jasmine.

### Short-read structural variant calling

Paired reads were first aligned to the *D. melanogaster* reference assembly (v6.58) with bwa-mem (Li 2013) via the “fq2bam” command within Parabricks (https://docs.nvidia.com/clara/parabricks/latest/index.html). Resulting bam files were merged with sambamba (Tarasov et al. 2015). Structural variants were called from short-read data using Manta (Chen et al. 2016), Delly2 (Rausch et al. 2012), Lumpy (Layer et al. 2014), and GRIDSS2 (Cameron et al. 2017). Variant callers were primarily used with default parameters. Manta’s break-end calls were converted to inversions with the “convertInversion.py” command. GRIDSS2 calls all variants as breakends, so the included script “gridssToBEDPE” was used to convert the calls to insertions, deletions, duplications, and inversions. Short-read structural variant calls were subjected to the same filtering steps as the long-read calls. Variants were required to be called by at least two callers when merging at the strain-level. Jasmine was used for all merging steps.

### Manual validation of structural variants

To verify the presence or absence of called structural variants in each of the inbred lines, we used Jbrowse2 (Diesh et al. 2023) to visualize the eight read alignments from the ultra-long read-length distribution. We elected not to visualize the master read pools as the extremely high coverages made loading all eight strains into visualization programs intractable. We required that the read alignments be in concordance with the VCF entries for multiple criteria for a structural variant to be positively verified. The read alignments had to show the same variant type, genomic location, and presence and absence across the eight lines as described in the VCF entry. When variants, specifically insertions, had small size differences between strains, we used UCSC’s Genome Browser (Perez et al. 2025) and the BLAT tool (Kent 2002) to ensure that all variants contained the same sequence. To maintain consistency, a single person performed all of the manual validation.

While inbred lines are generally treated as completely homozygous, there are still regions of heterozygosity (Frankham et al. 1993; Rumball et al. 1994; Powell and Evans 2017; De Kort et al. 2022) and we observed structural variants that were clearly heterozygous. We were able to differentiate heterozygous variants, even at low read frequencies, from sequencing errors or false alignment signatures. Heterozygous variants are often found in distinct haplotypes with different SNPs and indels in the surrounding genomic regions. All reads containing true heterozygous variants will similarly contain the same heterozygous SNPs and indels.

We briefly describe key alignment details for each type of structural variant. Non-tandem-duplication insertions should be found at the same breakpoint with the same length across the reads (Supplementary Figure 6). Tandem-duplication insertions should have the same length, but will not necessarily be found at the same breakpoint (Supplementary Figure 4). In shorter reads, duplications will be detected due to significant increases in coverage (Supplementary Figure 1). Depending on what the aligner deemed as the “duplicated” sequence, the breakpoint can range from the start of the first copy to the start of the second copy. Deletions should have the same length and same start and end breakpoints across reads (Supplementary Figure 7). Both insertions and deletions have specific graphical representations in JBrowse2, but in certain cases where the insertion or deletion is sufficiently large, it can be represented as a gap in aligned sequence. Inversions should have the same length and the same start and end breakpoints across reads. Small inversions that can be captured by single reads should clearly show the inverted sequence relative to the reference (Supplementary fig. 8). Large inversions will be characterized by split-mapped reads (Supplementary Figure 9). The reads on one side of a breakpoint should appear to map far away (in reference coordinate space) to the other end of the inverted sequence.

In the event of a false positive structural variant call, we classified the error into one of five broad categories. If the error stemmed from read misalignments of transposable or mobile elements or other repetitive elements as defined by RepeatMasker, then we classified the error as a “repetitive element” error (Supplementary Figure 2). If the variant error was caused by misalignments in a repetitive genomic locus (of non-repetitive-element sequence) or in a locus with intersecting or overlapping structural variants, the error was considered a “complex region” error (Supplementary Figure 1). If a variant call was found to be the summation of two smaller structural variants in the alignment, the smaller variants had differing sequence, and the region was not repetitive, then the call was considered a “strain merging” error (Supplementary Figure 5). If a putative variant call was found across multiple strains, and the variants in different strains were found to actually be different structural variants, then the putative call was assigned as a “population merging” error (Supplementary Figure 4). If a putative variant call did not correctly include a strain that actually had the variant, then the VCF was checked to see if the call was made at the strain level and incorrectly merged at the population level. If that was the case, then it was also designated as a population merging error. If the variant call did not exist at the strain level, then one of the other error designations was used. Finally, if a call did not fall into any of these categories, it was binned into “other” errors.

## Data access

All raw and processed sequencing data generated in this study have been submitted to submitted to the NCBI BioProject database (https://www.ncbi.nlm.nih.gov/bioproject/) under accession number PRJNA1247986. Raw Nanopore signal data (.pod5) will be provided upon email request due to their large file sizes. All genome assemblies and VCF files have been uploaded to Dryad (assemblies DOI: dx.doi.org/10.5061/dryad.8w9ghx3zj; VCFs DOI: dx.doi.org/10.5061/dryad.n5tb2rc6x). Detailed Snakemake pipelines for both genome assembly and structural variant calling are provided on github (https://github.com/jahemker/drosophila ultralong sv calling). A detailed DNA extraction and library prep protocol with our ultra-long-specific changes can be found on protocols.io (dx.doi.org/10.17504/protocols.io.14egn98wql5d/v1).

## Competing interest statement

The authors declare no competing interests.

## Acknowledgements

We thank the members of the Petrov lab and Tris Dodge for helpful comments and discussions about the manuscript. D.A.P was funded by the NIH NIGMS R35GM118165. D.A.P. is a Chan Zuckerberg Biohub – San Francisco Investigator.Some of the computing for this project was performed on the Sherlock cluster. We would like to thank Stanford University and the Stanford Research Computing Center for providing computational resources and support that contributed to these research results.

## Notes

### Competing Interest Statement

The authors have declared no competing interest.

