## Supplementary Information for "Manual validation finds only ultra-long long-read sequencing enables faithful, population-level structural variant calling in *Drosophila melanogaster* euchromatin"

### Supplementary Figures

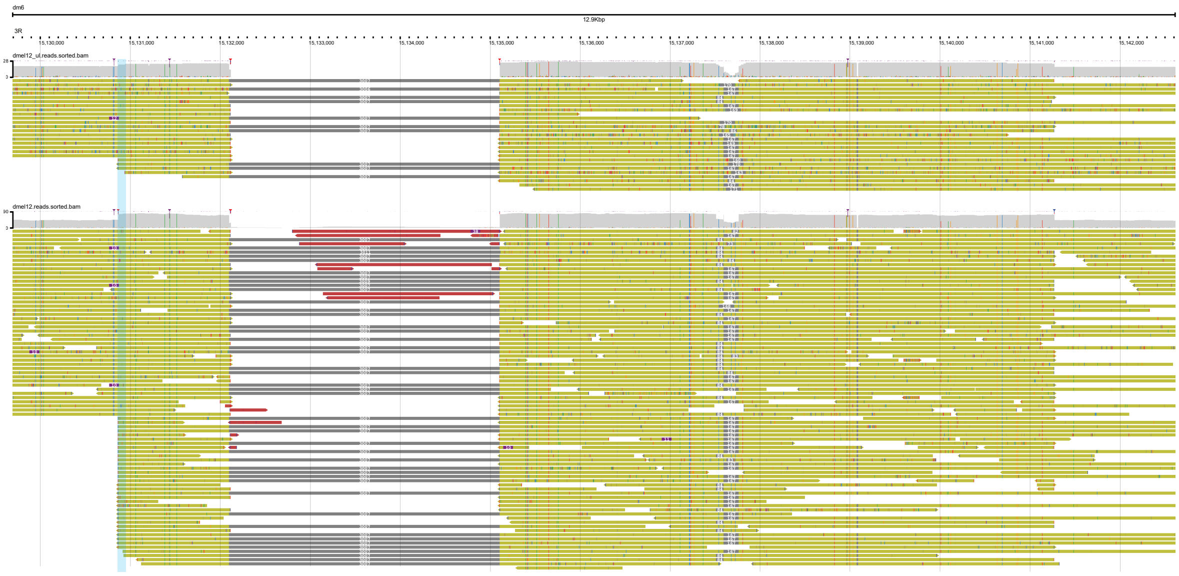


Supplementary Figure 1

**Supplementary Figure 1** An intersecting deletion and insertion were incorrectly called as a 10kb insertion, resulting in a “complex region” error. The blue highlighted region denotes the position of the putative variant call. The upper track is the ultra-long read alignment of the inbred line with the putative call. The lower track is the standard read alignment of the same line. Yellow reads have very high mapping quality scores while red reads have very low mapping quality scores. Reads that have red or blue ends are clipped. The coverage pileup shows this insertion is actually a duplication of a region that includes the genes *CG6912* and *CG3984*. Also clearly denoted in the reads is a 3kb deletion of a Jockey element. A duplication of a 10kb region containing a 3kb deletion should be called as a 7kb insertion.

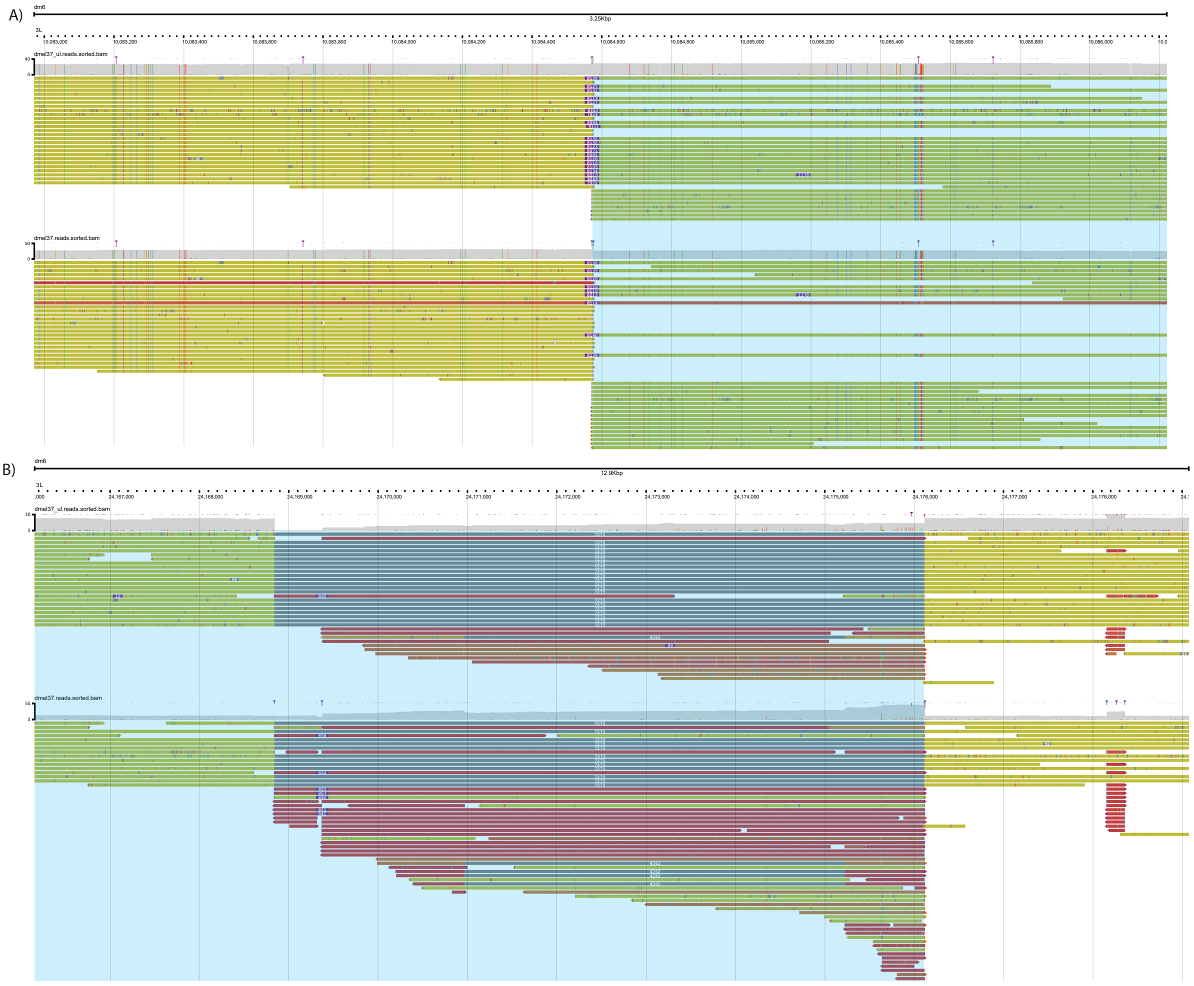


Supplementary Figure 2

**Supplementary Figure 2** A false inversion was called with the standard read-length data due to misaligned transposable element (TE) events. The blue highlighted region denotes the putative inverted sequence. The upper tracks show the ultralong read alignments, and the lower tracks show the standard read alignments. Yellow reads have very high mapping quality scores while red reads have very low mapping quality scores. A) The first putative breakpoint of the inversion call. The ultra-long read alignment clearly shows an 8kb insertion at this locus. A BLAT alignment of the inserted sequence finds that it is a Gypsy family TE. The standard read alignment has some reads that fully capture the insertion, but many more reads are clipped. These reads are not long enough for the aligner to properly place the insertion. B) The second putative breakpoint of the inversion call. The ultra-long reads clearly show a 7kb deletion. The deleted reference sequence is the same type of Gypsy TE that was inserted at the first breakpoint. In the standard read alignment, there are again a number of reads that cannot fully capture the deletion. Instead, the aligner has taken reads from the first breakpoint locus and mapped the TE portion to the second breakpoint locus, creating a false inversion signature.

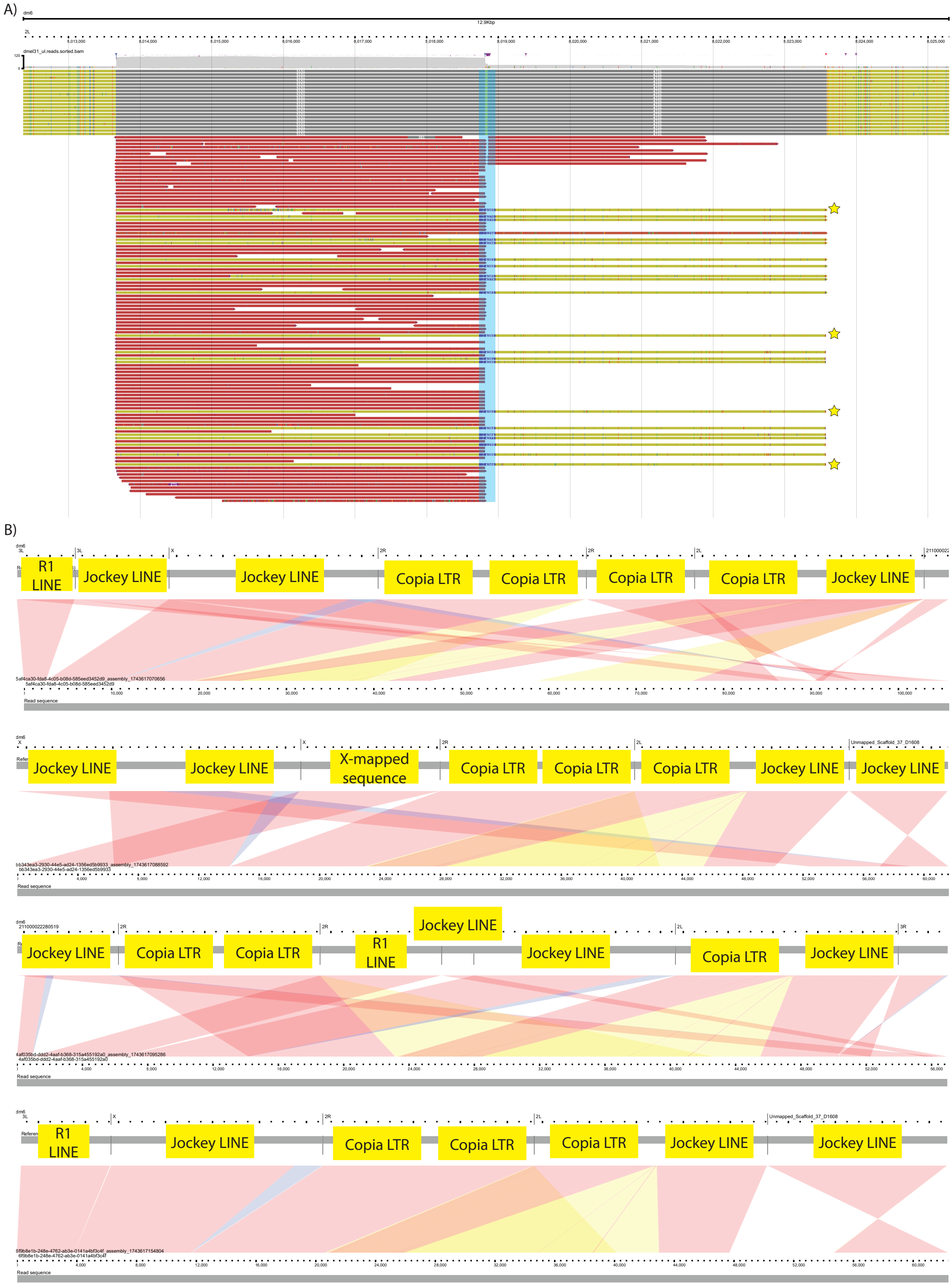


Supplementary Figure 3

**Supplementary Figure 3** Ultra-long reads incorrectly show a 16kb insertion due to read misalignment from an 80kb+ stretch of mobile elements. (A) The putative insertion locus is highlighted in blue. Yellow reads have very high mapping quality scores, while red reads have very low mapping quality scores. Most reads show two deletions of transposable elements in close proximity. We also see reads aligning to the deleted loci with a purple box in their centers, denoting the 16kb insertion. As the reads with the 16kb insertions do not map to the surrounding genomic region, it is unlikely that they are correctly mapped. B) We view four single-read alignments of reads that contain the 16kb insertion (starred reads in panel A). In each alignment, the reference sequence is at the top of the track, and the read sequence is at the bottom. The red bands connecting the reference and read sequences denote alignments. Yellow bands denote insertions, and blue bands denote deletions. Notably, all of the reads map to many loci on different reference chromosomes. We used BLAT to determine that nearly all of the sequences in these reads were different mobile elements, denoted by the yellow labels. These reads likely originated from a highly repetitive region of the genome, and the 16kb insertion is a false positive variant call.

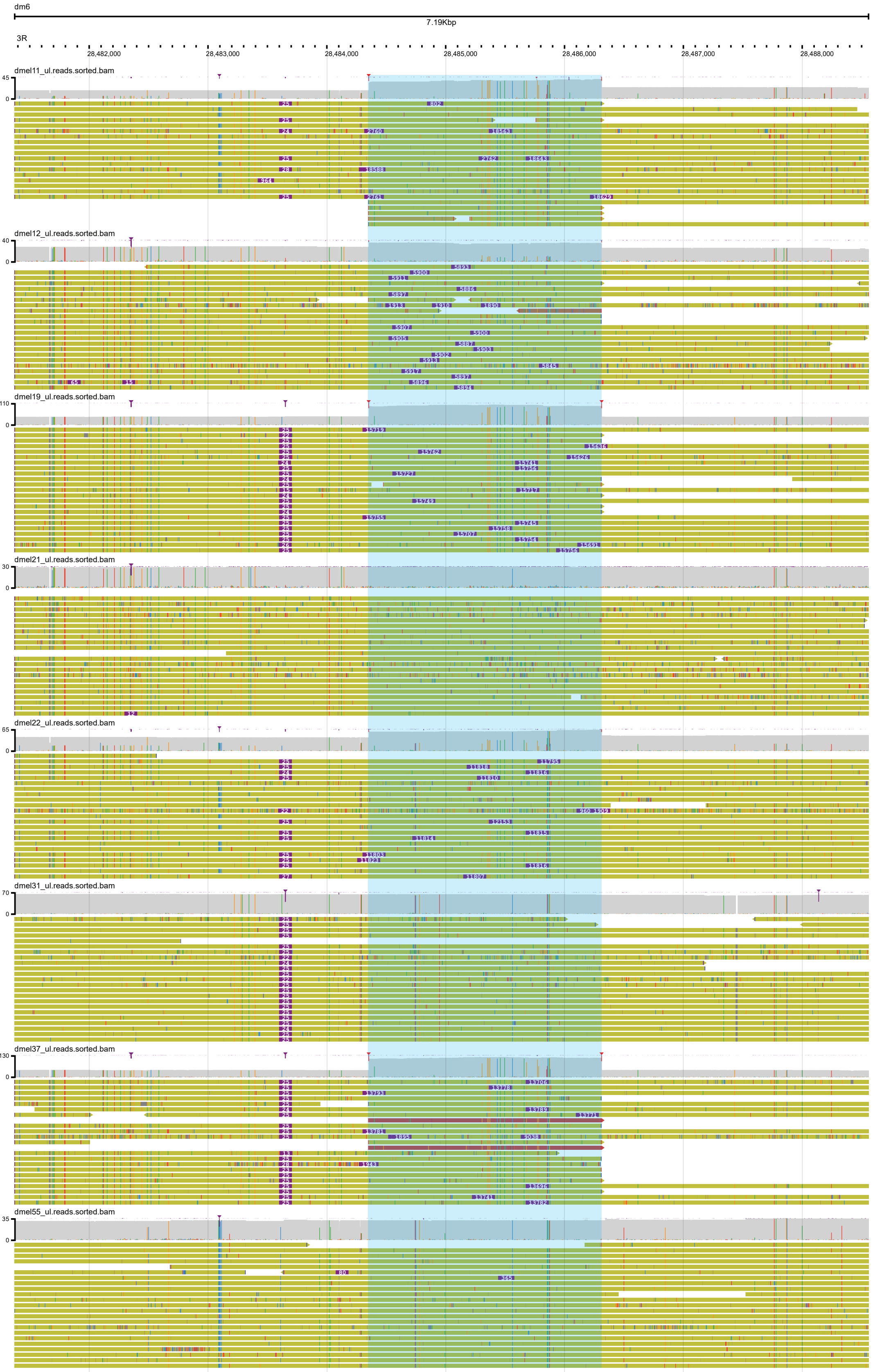


Supplementary Figure 4

**Supplementary Figure 4** A variable-number, tandem gene duplication was incorrectly merged into a single variant at the population level. The tracks are the ultra-long read alignments for each of the eight inbred strains. Yellow reads have very high mapping quality scores, while red reads have very low mapping quality scores. The genomic region highlighted in blue is the gene *Myst5*. The grey track above each read alignment denotes read coverage. Corresponding with the coverage increases are insertions, ranging in size from 6kb-18kb across five different lines. BLAT alignments from each of the strains found that the insertion sequences were tandem duplications of *Myst5*. The variation in length stemmed from different copy numbers of *Myst5*. The insertions varied in breakpoint, both between and within strains, but all were placed within the *Myst5* locus which was expected for a tandom duplication alignment. The putative variant call was a 13kb insertion found in the lines dmel12, dmel21, dmel22, dmel31, and dmel37. While there were insertions in each of these lines, only dmel37 had an insertion close to size 13kb. The copy number variation found across the lines was incorrectly lost when all of these different duplications were merged together at the population level.

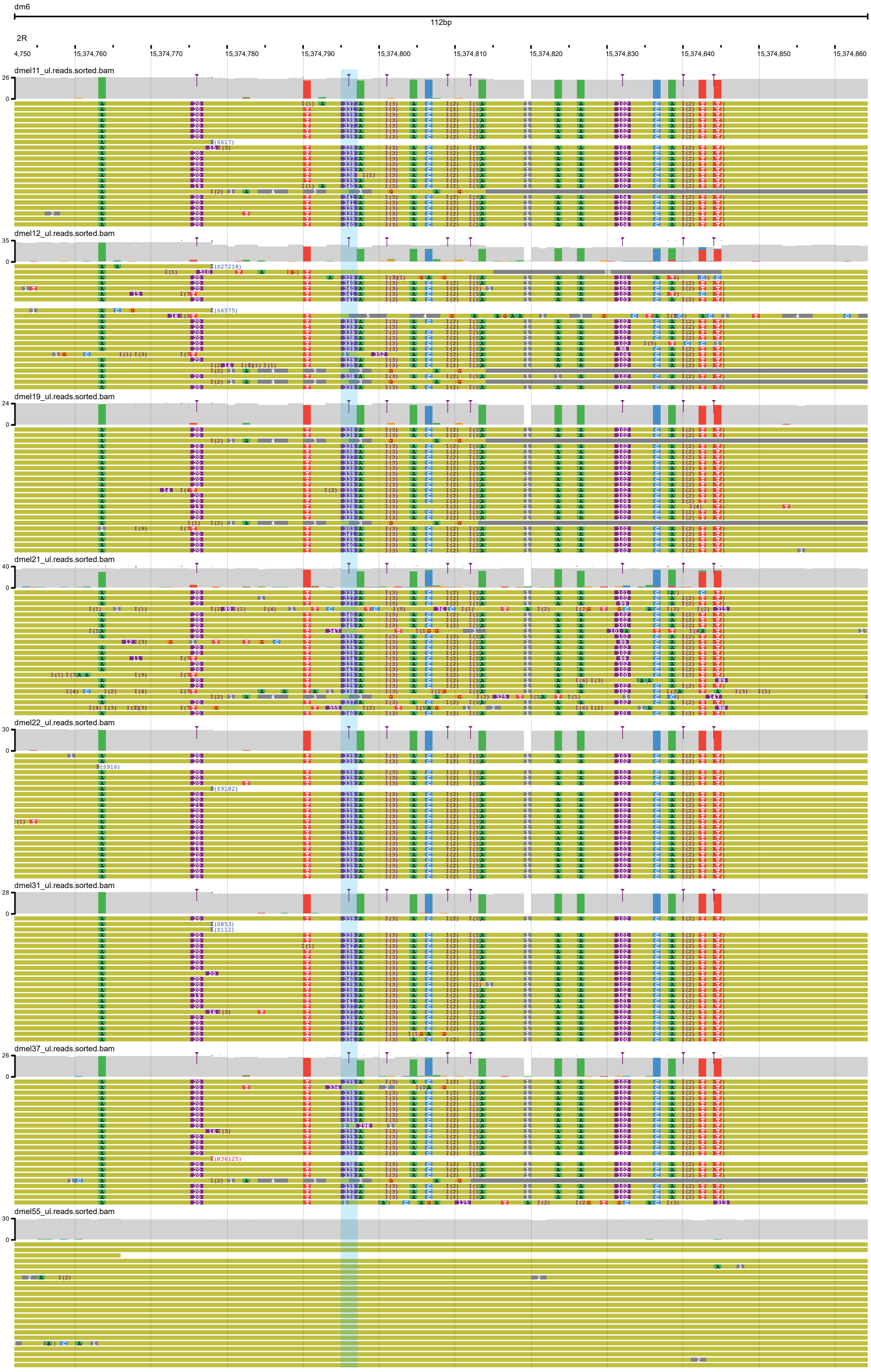


Supplementary Figure 5

**Supplementary Figure 5** Two insertions were incorrectly merged into a single insertion at the strain level. The tracks are the ultra-long read alignments for each of the eight inbred strains. Yellow reads have very high mapping quality scores. The putative variant call denoted a single 440bp insertion at the blue, highlighted locus in the seven lines. The read alignments clearly showed seven inbred lines with three insertions of lengths 20bp, 100bp, and 340bp, suggesting that the callers merged the 340bp insertion with the 100bp insertion. Investigating the specific inserted sequences found that the 340bp belonged to the PROTOP_A repeat family, while the 100bp belonged to the similar, but distinct, PROTOP_B repeat family.

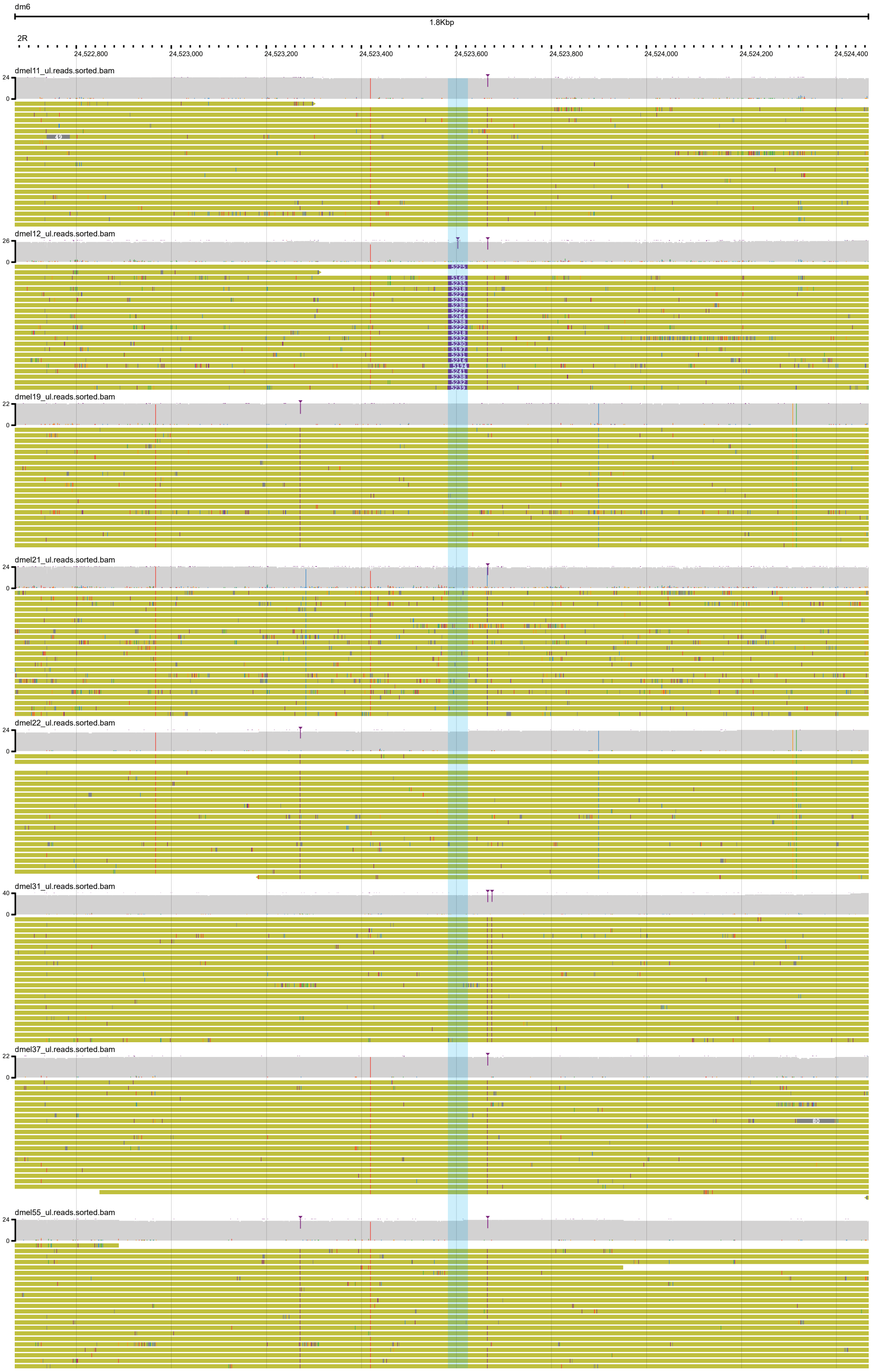


Supplementary Figure 6

**Supplementary Figure 6** A true insertion call. Each of the eight tracks shown are the ultra-long read alignments for the inbred lines. Yellow reads have very high mapping quality scores. The blue highlighted region marks the putative insertion breakpoint. This putative variant call denoted a 5.2kb insertion only found in dmel12, which can be clearly observed.

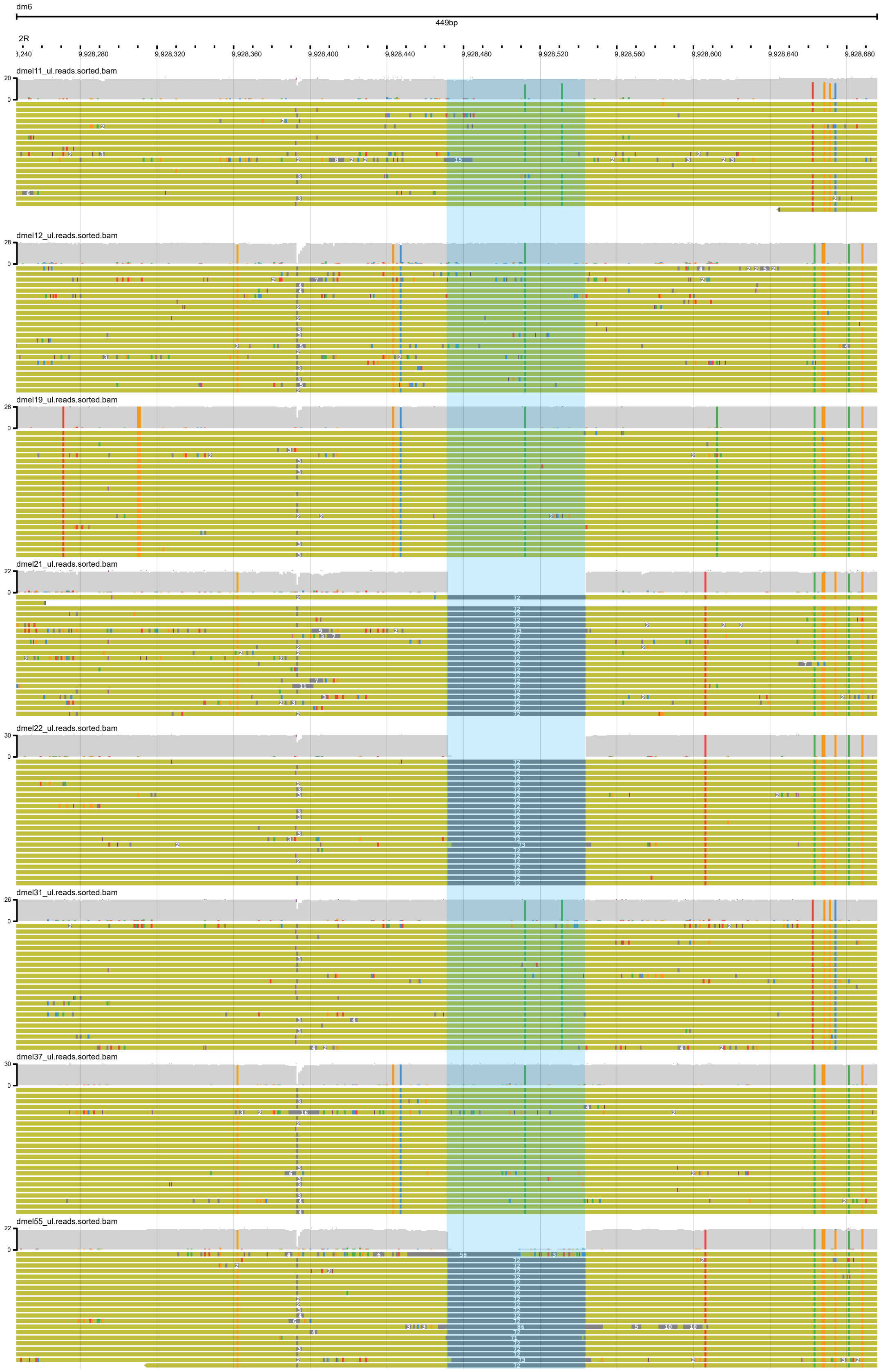


Supplementary Figure 7

**Supplementary Figure 7** A true deletion call. Each of the eight tracks shown are the ultra-long read alignments for the inbred lines. Yellow reads have very high mapping quality scores. The blue highlighted region marks the putative deletion locus. This putative variant call denoted a 72bp deletion in dmel21, dmel22, and dmel55, which can all be clearly observed.

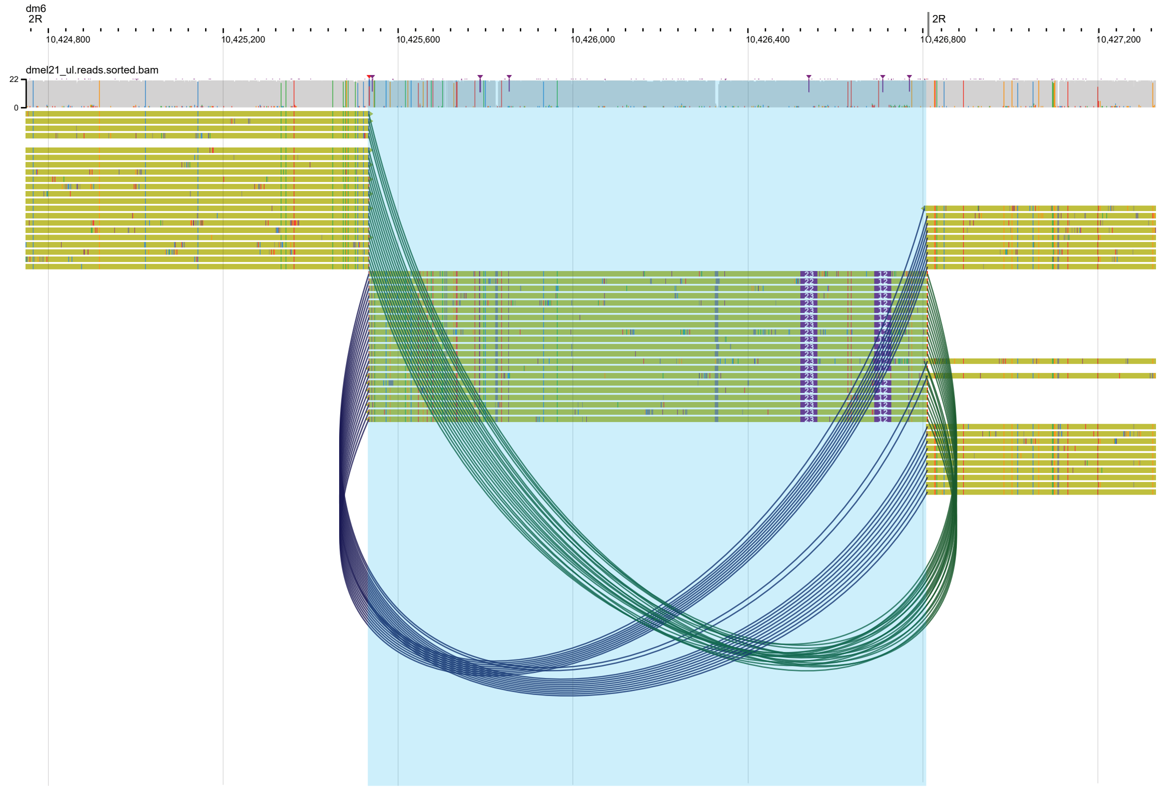


Supplementary Figure 8

**Supplementary Figure 8** A true, small inversion call. The track is the ultra-long read alignment for the strain in which the putative inversion was called. Yellow reads have very high mapping quality scores. This putative variant call denoted an inversion of size 1.2kb, highlighted in blue. This is small enough such that the full inversion should be captured in the individual reads. Split-mapped read alignments are denoted with green and blue connecting lines. Following the genomic path shown by the connecting lines, it is clear in all reads that the central 1.2kb sequence is actually an inversion.

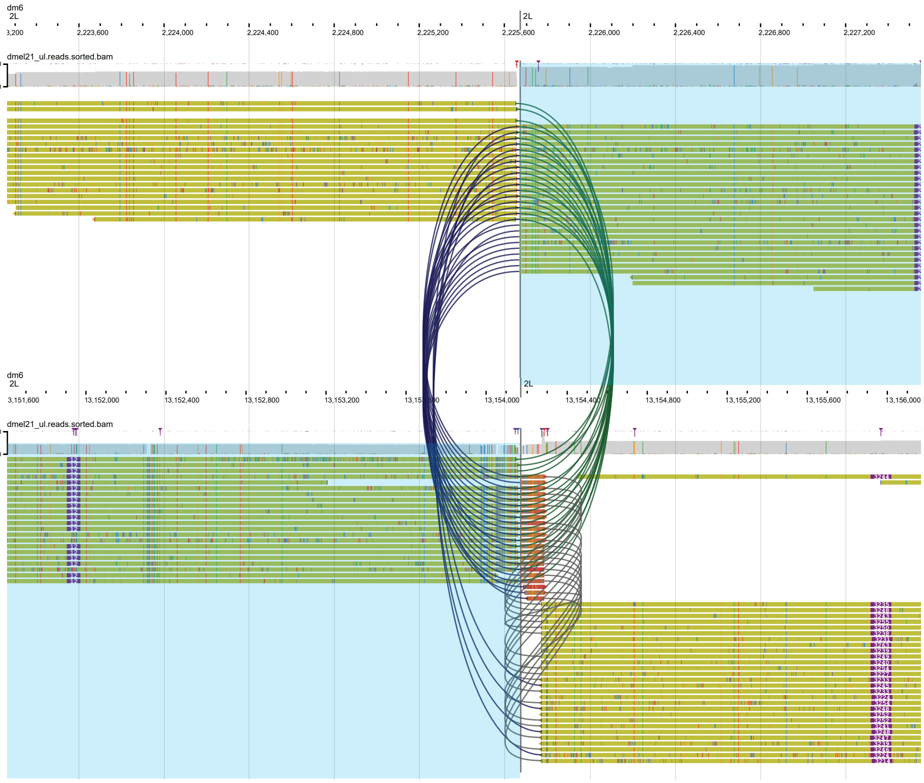


Supplementary Figure 9

**Supplementary Figure 9** A true, large inversion call. Both tracks are the ultra-long read alignment for the strain in which the inversion was called, with each track focusing on a putative breakpoint locus. Yellow reads have very high mapping quality scores. The blue highlighted region denotes the putative inversion locus. Split-mapped read alignments are denoted with green, blue, and gray connecting lines. This putative variant call denoted an inversion of size 10.9Mb. Reads at the inversion breakpoints should show clear mapping to either end of the inversion. As expected, the reads before the first breakpoint at 2L:2,224,600 show alignment to the inversion end at 2L:13,154,000. The same is true for the other breakpoint. This is a true inversion call of the cosmopolitan inversion In(2L)t.

### Supplementary Tables

**Supplementary Table 1**
Sequencing and assembly statistics per inbred line.

*(See excel spreadsheet.)*

| **Chromosome** | **Start** | **End** |
|---|---|---|
| 2L | 82455 | 19570000 |
| 2R | 8860000 | 24684540 |
| 3L | 158639 | 18438500 |
| 3R | 9497000 | 31845060 |
| X | 277911 | 18930000 |

**Supplementary Table 2** Euchromatic regions for each chromosome, following Chakraborty et al. (2018).

| Berhmann et al. 2015 Line ID | Hemker et al. 2025 Line ID |
|---|---|
| 12LN1057 | dmel11 |
| 12LN1059 | dmel12 |
| 12LN1083 | dmel19 |
| 12LN1086 | dmel21 |
| 12LN1089 | dmel22 |
| 12LN630_(B66) | dmel31 |
| 12LN649_(B9) | dmel37 |
| LNPA36 | dmel55 |

**Supplementary Table 3** List of *D. melanogaster* inbred lines used in this study from Behrman et al. (2015).
